# Substrate mediated elastic coupling between motile cells modulates inter–cell interactions and enhances cell–cell contact

**DOI:** 10.1101/2021.03.06.434234

**Authors:** Subhaya Bose, Kinjal Dasbiswas, Arvind Gopinath

## Abstract

The mechanical micro–environment of cells and tissues influences key aspects of cell structure and function including cell motility. For proper tissue development, cells need to migrate, interact with other neighbouring cells and form contacts, each of which require the cell to exert physical forces. Cells are known to exert contractile forces on underlying soft substrates. These stresses result in substrate deformation that can affect migratory behavior of cells as well as provide an avenue for cells to sense each other and coordinate their motion. The role of substrate mechanics, particularly its stiffness, in such biological processesis therefore a subject of active investigation. Recent progress in experimental techniques have enabled key insights into pairwise mechanical interactions that control cell motility when they move on compliant soft substrates. Analysis and modeling of such systemsis however still in its nascent stages. Motivated by the role modeling is expected to play in interpreting, informing and guiding experiments, we build a biophysical model for cell migration and cell–cell interactions. Our focus is on situations highly relevant to tissue engineering and regenerative medicine –when substrate traction stresses induced by motile cells enable substrate deformation and serve as a medium of communication. Using a generalizable agent–basedmodel, we compute key metrics of cell motile behavior such as the number of cell–cell contacts over a given time, dispersion of cell trajectories, and probability of permanent cell contact, and analyze how these depend on a cell motility parameter and on substrate stiffness. Our results provide a framework towards modeling the manner in which cells may sense each other mechanically via the substrate and use this information to generate coordinated movements across much longer length scales. Our results also provide a foundation to analyze experiments on the phenomenon known as durotaxis where single cells move preferentially towards regions of high stiffness on patterned substrates.

## 1. Introduction

Many eukaryotic cells move by crawling, that is by adhering to and exerting mechanical stresses and local forces on their extracellular matrix (ECM) that they then actively deform (see for instance [1–4] and references therein). Existing approaches to modeling collective cell motility focus on direct (steric and adhesive) cell-cell interactions or focus at the single cell level on cell-substrate interactions [2] such as the details of focal adhesions that are crucial to generating traction stresses in both adherent and motile cells [5]. Experiments strongly indicate however that cells cultured on soft, elastic, biocompatible substrates can respond to each other even when not in direct contact [3,4]. Such interactions can arise in cell culture experiments, with cells on the surface of synthetic hydrogels such as polyacrylamide, which are linearly elastic, through mutual and active deformations of the gel by the cells. These mechanically derived non-contact cell-cell interactions are even more relevant and act over longer ranges in the biological extracellular matrix (ECM) comprising collagen or fibrin, where cells can interact by remodeling and reorienting the fibers in the ECM [6–8]. Even without such cell–matrix feedback, the presence of deformations has been shown recently to guide the migration of other cells without requiring chemotactic cues [9].

Mechanical non-local interactions between cells offer advantages compared to chemical means. Specifically, mechanical signaling and mechanosensing of neighbouring cells is typically faster and longer-ranged than chemical signaling. Chemical interactions are limited by diffusion rates while mechanical interactions propagate near instantaneously for purely elastic deformations [10]. Indeed, this crucially allows cells to not just sense each other but also to synchronize their behaviour. For instance, substrate deformation-mediated long-range interactions has been clearly demonstrated in heart muscle cells that synchronize their beating without direct contact [11,12], as well as at a subcellular level between myofibrils within a single heart muscle cell [13]. Cell communications via sensing of substrate or matrix deformation are particularly important in sparse, non-confluent cell cultures or tissue that occur in a number of biologically relevant situations. Apart from beating cardiomyocytes, examples of such situations include wound healing involving fibroblasts [14], sprouting blood vessels comprising endothelial cells [15], and migration of mesenchymal cells in zebrafish embryo before the formation of confluent epithelial tissue [16]. In all these cases, cells are not in direct contact but exert traction forces on the surrounding mechanical medium and concomitantly sense deformations caused by nearby cells. Such interactions therefore crucially depend on the stiffness of the substrate, and can be probed by experiments that vary the stiffness of the hydrogel substrate on which the cells are cultured [17,18]. These aspects influence not only motility response at the single cell level but also strongly impact collective behavior including directed motility and subsequent spatial self-organization.

On the other hand, while substrate-mediated cell-cell elastic interactions have been considered for the organization of adherent cells in a variety of mechanobiological contexts [19,20] (the physical basis of such modeling is reviewed in Ref. [21]), their effect on collective cell motility, which in principle is always present, have not been carefully modeled. Here, we present a simple biophysical agent–based model and computational results that focus on how substrate mediate mechanical communication allows two cells to sense each other and impacts their collective and relative motility. Our approach provides a foundation for the study of more general cell interactions that include both mechanical and chemical signalling, and also serves as a starting point for future studies of mechanical substrate based interactions in multi-cellular systems such as growing tissue and confluent sheets.

## 2. Experimental observations motivate model for cell elastic interactions

Many eukaryotic cells use contractile localized forces generated by their actomyosin cytoskeleton to adhere to and move on their substrates. Such traction forces typically cause measurable deformations in the underlying substrates in cell culture experiment [5], and have a spatially dipolar pattern [22]. A cell typically acts as a force dipole exerting – a pair of equal and opposite forces – on the elastic medium. The dipolar pattern arises due to the fact that no external forces are present on the system, and the cell, as a whole, moves on its own accord. The net effect of these stresses is to contract or pull in the elastic material comprising the substrate towards the cell.

In Ref. [3], it was shown that endothelial cells cultured on hydrogel substrates of varying stiffness change their motile behavior in the presence of traction stresses exerted on the substrate by neighbouring but non-contacting cells. In particular, it was shown that pairs of cells on softer gels, showed reduced collective migration in comparison to isolated cells. The number of contacts two cells made over specific periods of time by extending their pseudopodia towards each other was also measured and found to depend sensitively on substrate stiffness. Remarkably, the cells made stable contacts on very soft gels (Elastic modulus, 500 Pa), whereas they made repeated contacts and withdrawals on substrates of intermediate compliance (Elastic modulus, 2500 – 5500 Pa).

Motivated by this experiment, we here model the motility characteristics of a two cell system and demonstrate how elastic deformations induced in the substrate allow cells to respond to each other. We consider a pair of cells that each adhere to, and exert stresses on the underlying substrate thereby deforming it as shown in Fig. 1. As mentioned above, adherent and motile cells generate a contractile stress on the substrate. Here, the contractility **P** of each cell, is minimally described by a physical model of force dipoles –a pair of equal and opposite forces exerted on the substrate, and is thus a tensorial quantity [19]. Such modeling is inspired by the theory of deformations induced by inclusions in materials [23]. Unlike passive material inclusions, cells can actively regulate their force production in response to external mechano-chemical cues from the substrate, including the presence of other cells. Such complicating feedback effects in cell–cell interactions has also been theoretically considered [24,25], but we ignore these for simplicity here, and we treat *P* as an intrinsic cell property that is independent of underlying substrate matrix strain and stiffness.

**Figure 1.**
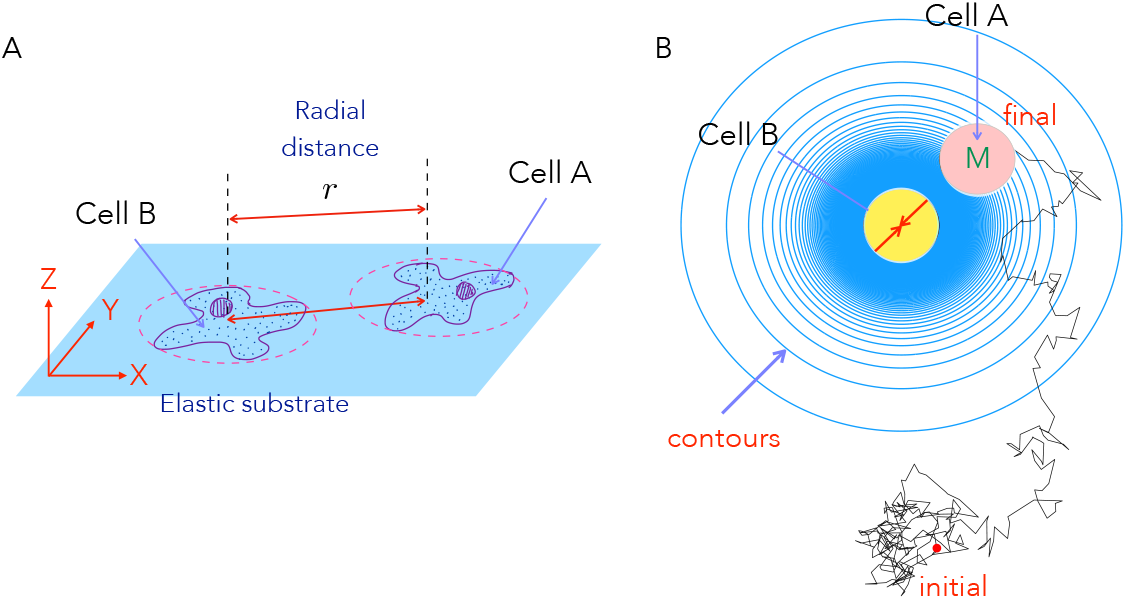
Schematic of the cell-cell mechanical interactions model: (A) Two cells A and B cultured on the surface of thick elastic substrate can sense each other and interact at long range (when the inter-cell distance *r* is longer than typical cell sizes, here depicted by dashed red circles) through mechanical deformations of the underlying substrate. These deformations arise as the cells exert traction stresses on the underlying elastic material. Here the cells are restricted to move on the surface of the substrate. (B) We study with our computational model how a motile cell (M, Cell A, pink) moves in the presence of a fixed central cell (Cell B, yellow). This two cell system on a substrate (schematic shown as a top view) also mimics scenarios where a motile cell may encounter an elastic impurity or obstacle on the medium. Shown as blue circles are contours of constant elastic potential (in simplified form) that determine the inter-cell elastic force experienced by the motile cell B as a result of the elastic deformations of the medium by both cells A and B. Also shown (in black) is a representative simulated trajectory of the motile cell which starts outside the *area of influence* of the stationary cell.

To simplify our study, we assume that one of the cells is motile (Cell A) and the other is stationary (Cell B). The stationary cell B is nonetheless *alive* in that it still deforms the substrate. The resulting deformation field, or equivalently the substrate mediated elastic potential, is sensed by the other, distant, motile cell A. The interaction potential between the cells in turn creates a mechanical force on the motile cell A. For polarized and elongated cells, the deformations have a dipolar spatial pattern (described in Appendix A). However, here we consider a simplified scenario that is valid when cells reorient very fast in the time for them to translate and migrate (Appendix A, §3). This implies that the directions of the dipole axis (of both cells A and B) fluctuates rapidly as cell A moves resulting in an effectively isotropic, attractive interaction potential that decays with distance as ~ 1/*r*^3^ (iso–surfaces shown as blue circles in Fig. 1 B). Analysis of this model interaction provides us insight into attractive potentials strongly influence cell motility.

The motile cell is considered to move diffusively with an effective diffusion coefficient, while also being acted upon by an elastic interaction force from the stationary cell. Although, polarized cells may propel themselves persistently along their body axis, we consider more isotropic cells here which extend their pseudopodia in different directions randomly, and are thus described adequately by a diffusive process. Such a simple effective Langevin equation is commonly used to describe elastically coupled motile active particles [26] and swarming bacteria [27] but has not been studied previously in conjunction with this specific type of interactions that arise on an elastic substrate.

We note that the model can be easily generalized (as derived in Appendix A) to describe a pair of motile cells since the interactions are pairwise and reciprocal. The interaction potential is not isotropic and depends on both the inter-cell distance as well as on the instantaneous alignment of the cells’ dipole axes. Thus the force on each cell (related to the gradient of the potential) depends on not just the relative positions of the cells.but additionally on the direction of the contractile dipoles exerted by cells A and B. Truly spherical dipoles embedded in an elastic medium do not interact mechanically [23], unless cell-substrate feedback effects occur [25]. Furthermore, cell-cell interactions in a fibrous, nonlinear elastic medium can be longer ranged [28] and have a power law character, ~ 1/*r^α^*, where *α* < 3 [29]. The interaction of disk-like cells on top of a thick substrate (semi-infinite geometry) is also more complicated [30]. We choose the isotropic, attractive 1/*r*^3^ potential as the simplest attractive interaction with the same distance dependence as the dipolar interaction, with the objective of testing how such a potential can affect cell motility. Motivating future work, we show how the conclusions from the simpler potential remain qualitatively valid even as specifics of cell trajectories change when the more general dipolar potential is used. This model highlighted in this work, although very simplified both in its description of cell contractility and motility, can thus capture key aspects of motility and contact formation, as we now describe.

## 3. Materials and Methods

### 3.1. Model for two-cell interactions

The model used to analyze the two-cell system is an agent-based stochastic model. We start with the stochastic Langevin equation for the dynamics of the moving cell *A* in the presence of a second cell *B* fixed at the origin as illustrated in Figure 1(A). Details of the model and the simplifications involved may be found in Appendix A. Starting from the more general model where both cells A and B can move, we now fix cell B and thus set **r^*B*^** = **0**. In other words, we choose the center of cell B to be the origin from which the position of cell A and its distance relative to B is measured. Writing **r** = **r^*B*^** – **r^*A*^**, we write the equation for **r**(*t*) where *t* is the time,

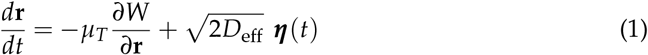

where *D*_eff_ is the effective translational diffusivity quantifying the random motion of the moving cell in the absence of the fixed cell, and ***η*** is a random white noise term whose components satisfy

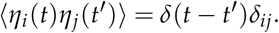

Note that *η* - the active noise term - has units of *t*^−1/2^. The mobility *μ_T_* in equation (1) quantifies the effective friction from the medium and is inversely proportional to the cell size *σ* and inversely proportional the the viscosity at the surface. Here it is assumed that the cells moving on a wet surface and that the fluid nature of the surface provides a viscous resistance opposing cell motion.

The two-cell potential W derives from the elastic interactions communicated via the linear deformation of the substrate (Appendix A, Equation A5) and is given by,

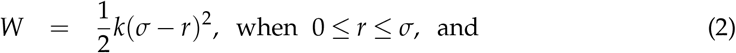

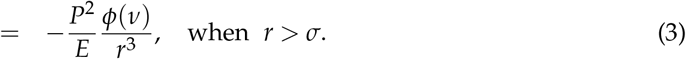

Numerical solutions to equation (1) are obtained with varying initial conditions for cell *A* as explained subsequently. To ease the computational analysis, we work in scaled dimensionless units. We choose cell size (diameter) *σ* (see Fig 1), diffusion time *σ*^2^/*D*_0_, and thermal energy *k_B_T* – with *T* corresponding to the temperature of the cell/substrate system – as our length, time and energy scales respectively. Equations (1–3) may then be rewritten as

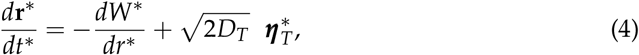

where the potential in scaled form is

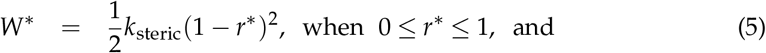

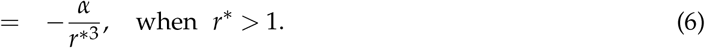

Superscripts * in equations (4)–(6) denote non-dimensional quantities. Henceforth, we will drop this subscript for clarity. Thus the dynamics may be followed as a function of three dimensionless numbers (parameters)

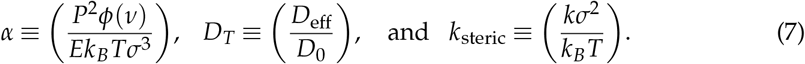

### 3.2. Dimensionless parameters quantifying cell motion and interactions

The parameters that emerge in equations (1)–(7) and typical of the two-cell scenario studied here are summarized in Table 1. Following Ref. [3], we are interested in substrates that are linearly elastic with the Young’s modulus *E* ranging from 0.5 kPa to 33 kPa, well within the range of 0.1-100 kPa appropriate for tissues and bio-compatible materials [18]. The effective diffusion coefficients exhibited by cells in experiments [3] include the random noisy motion as the cells explore territory and a contribution due to short-time deterministic motion. We explore values in the range 3*μ*m^2^/minute to 50 *μ*m^2^/minute. Time scales are estimated from experiments as well and 250 seconds in real time correspond to a dimensionless time duration of unity.

**Table 1.** Biophysical parameters characterizing the two-cell (typical values from [3,31,32]).

| Quantity | Interpretation | Experimental values |
| --- | --- | --- |
| $\sigma$ | Cell size | 10-100 $\mu\text{m}$ |
| $T$ | Temperature | 25<sup>0</sup> C |
| $D_0$ | Thermal Diffusivity | 25 $\mu\text{m}^2/\text{min}$ |
| $D_{\text{eff}}$ | Effective Diffusivity | 3 – 50 $\mu\text{m}^2/\text{min}$ |
| $E$ | Young's modulus | 0.5 – 33 kPa |
| $\nu$ | Poisson ratio | 0.3 - 0.5 |
| $P$ | Contractility | 10<sup>-14</sup> Nm |

Scaled non-dimensional parameters relevant to the simulation may be calculated from dimensional quantities as explained earlier. Three scaled parameters determine the dynamics of the two-cell system: *D_T_*, *α* and *k*_steric_. Values used in the computations are listed in Table 2. The self avoidance parameter *k*_steric_ is chosen such that the cells don’t overlap and is computed based on the time step used in the simulations. This allows us to control the stability of the simulation and its accuracy.

**Table 2.** Simulation parameters and their meaning.

| Parameter | Interpretation | Definition | Simulation values |
| --- | --- | --- | --- |
| $D_T$ | Diffusivity | $D_{\text{eff}}/D_0$ | 0.1-10 |
| $\alpha$ | Cell-cell interaction | $P^2\phi(\nu)/(Ek_B T\sigma^3)$ | 0.1-100 |
| $k_{\text{steric}}$ | Self-avoidance | $k\sigma^2/k_B T$ | 10<sup>3</sup> – 10<sup>4</sup> |

### 3.3. Numerical solution and tracking cell trajectories

Equations (4)–(7) are solved for the dynamics of the moving cell with appropriate boundary and initial conditions. The Langevin equation (4) is an example of stochastic differential equations; here we solve this equation using the explicit half-order Euler-Maruyama method one of us has used recently in similar problems involving bacteria cells moving in light fields [27] and in simulations of active Brownian particles [26].

Given the position of cell A at time *t*, **r**(*t*), its subsequent location at time *t* + *δt*, **r**(*t* + *δt*), follows,

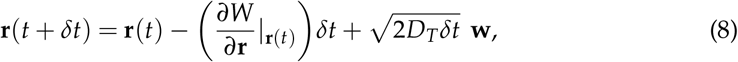

where **w** is a random two-dimensional vector with components each drawn at every time step from a normal distribution with mean zero and standard deviation of unity.

We simulated several trajectories of cell A ((*n* = 1000) trajectories, diameter *σ* = 1 in scaled units), under the influence of the central stationary cell B (also having diameter *σ* = 1). The simulations were conducted in two different geometries as described below.

To study the contact frequency between two-cells and explore the systematically explore the role of the elastic potential, we simulated cell A moving in a confined square box of size 12*σ* with the stationary cell B at the center of the box. Cells reflect from the box surface when they encounter it and thus are restricted to remain within the simulation domain.

In order to calculate the number of contact in due course of the simulation, we define a contact radius 1.5*σ* from the centre of the stationary cell, and we consider a contact if the centre of the test cell lies within the contact radius. The cell can come out of the contact radius and re-enter, increasing the number of contacts. The time step used in these simulations is *dt* = 0.0001 and total number of steps in this simulation is 10^7^, i.e. a cell trajectories were followed for a total time of *T* = 1000.

On the other hand for calculating cell dispersivities, and specifically the mean squared displacement (MSD) of cell A, we used periodic boundary conditions and a periodic potential. This corresponds to cell A moving in a periodic domain and interacting with a regular square lattice of multiple stationary cells (images of B) separated uniformly by a distance 12*σ*. The time step used to integrate equation (7) in these simulations is also *dt* = 0.0001 and total number of steps in this simulation is 10^7^, i.e. a cell trajectories were followed for a total time of *T* = 1000. The mean square displacement MSD was calculated by tracking trajectories of cell A (the same as tracking *n* = 100 cells). As before, cell A is initialized randomly inside the same square box of length of 12*σ*, but outside the contact radius. Cells that move out of the domain are reintroduced into the domain in a manner that respects periodic boundary conditions and the appropriate symmetries.

In this case since **r** ≡ *x***e**_*x*_ + *y***e**_*y*_ is the relative distance between the cells, the mean square displacement is calculated by the equation,

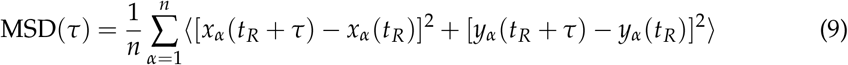

where *τ* is the delay time, and the summation is over each cell trajectory (indexed by *α*) and extends over the full number of trajectories *n* = 100. The delay time is varied and the averages are obtained by choosing different values of the reference time *t_R_* as is normally done. The MSD given by equation (9) is thus an average over time and also an average over realized cell trajectories.

The mobility of cell A reflects the properties of the microenvironment created by cell B and by the substrate. The mean square displacement in (9) is written as a function of the delay time *τ* that may be interpreted as an effective observation time over which the cell motion is observed. For instance, a cell that moves with constant speed for small times (say ~ *T*_1_) and undergoes a diffusive random walk when observed over long times (say ~ *T*_2_) will exhibit different slopes for *τ* < *T*_1_ and for *τ* > *T*_2_. The exponent characterizing the dependence of the MSD on the delay time provides information as to whether the motion is sub-diffusive (exponent < 1), diffusive (exponent = 1), or super-diffusive (exponent > 1).

It is constructive to study the expected MSD for cell A in the absence of cell B. In this particular case, since A is purely diffusive, the MSD has the simple form valid for diffusion in two dimensions MSD(*τ*) = 4*D_T_τ*. Deviations from this expression arise due to the mechanically induced inter-cell interaction and thus quantify the extent to which cell B perturbs the dispersion of cell A. For instance transient or persistent trapping of cell A will result in the MSD scaling sub-linearly with *τ*.

## 4. Results

### 4.1. Cell-cell contact frequency shows biphasic dependence on matrix elastic interactions

Motivated by experiments which show that two cells make repeated contact and withdrawals on soft substrates, with contact frequency dependent on the substrate stiffness, we measure the total number of contacts of the motile cell (A) with the stationary cell (B) in our model simulations. As indicated earlier, the simulated cells are initialized randomly inside the box, but outside of a pre-defined contact radius around the stationary cell. The total number of contacts between the cells is counted over a fixed period of time i.e. T = 1000. It should be remembered that the cells are confined to stay within the square domain during the course of the simulation.

Cell A’s movement is governed by an attractive elastic potential induced by the stationary, central cell and its own random motion, described as an effective diffusion. Additionally when the cell encounters the bounding wall of the square domain, it reflects (moves away) from it. Overall, random noise encapsulated in the diffusion coefficient causes A to move towards or away from B in an unbiased manner. The attractive potential *W* being isotropic and spatially varying suggests that there is a critical radius of influence (dependent on both *α* and *D_T_*) within which forces due to the attractive potential dominate diffusion and significantly influence the trajectory of cell A. This effect results in the cell getting closer to cell B, eventually entering this zone of influence.

To carefully study how elastic interactions (*α*) and random diffusion (*D_T_*) each influence this process, we first systematically calculated the number of contacts by *α*, while keeping *D_T_* constant at three different values, *D_T_* = 1, 2,5. (Figure 2). As illustrated by the dotted lines which serve as a guide to the eye, the behavior is highly non-monotonic. For small *α*, the number of contacts increases with increasing *α*, then reduces to 1 at high *α*. The position of the peak increases with increasing *D_T_*. The initial increase in contacts is due to the increased directional movement of the test cells towards the central cell. The decrease in the number of contacts for very high values of *α* is expected since the attractive potential is strong enough to overcome the effect of diffusion. In this case, the motile cell is unable to move away from and makes stable contact with the stationary cell. For *α* = 5 and *D_T_* = 1 (trajectory 1), the test cell spends most of the time exploring space rather than near the stationary cell, which also reduces the number of contacts. Increasing *α* to 10 (trajectory 2) the radius of influence increases, increasing the duration of contact and thereby increasing contacts. On further increasing *α* to 20 (trajectory 3), the test cell is tightly adhered to the stationary cell which allows only one single contact. Note that the statistics for the high *D_T_* and low *α* regime are influenced by the confinement. Cells in this particular limit frequently escape the region of influence and wander away only to return again after encountering the wall and diffusing away. For instance, the number of contacts for *D_T_* = 5 and *α* = 0.1, combines the effect of repeated escapes from the region of influence and repeated returns due to confinement. Since the size of the box is fixed, the increase in number of contacts with *D_T_* for *α* = 0.1 is still a signature of diffusive effects dominating the attractive potential.

**Figure 2.**
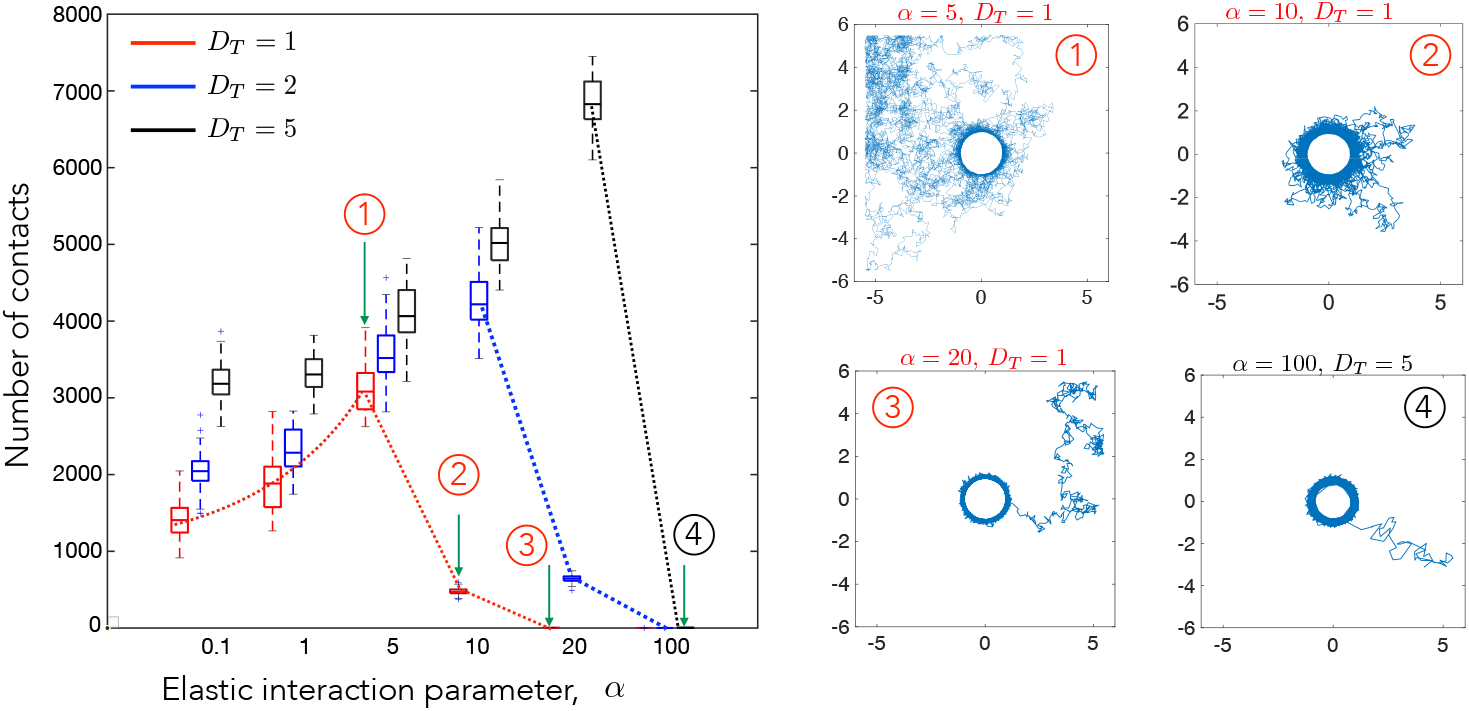
The number of cell–cell contact events measured in a fixed interval of time depends strongly on the elastic interaction parameter. A contact event is identified as cell A coming within a prescribed *contact radius* of cell B with cell A initialized randomly in a certain area around cell B. Thus the number of contact is be interpreted as the average number of contacts of the two cells. The number of simulation runs conducted were 50 for each combination of *D_T_* and *α*. The dashed curves are guides to the eye illustrating the trends seen with increasing values of *α*. Diffusion is the major factor in governing the number of contacts for low values of *α*. For higher *α*, the attractive potential increases the probability of the cell to stay near the contact radius and controls the number of contacts. Trajectories for highlighted data points (1)-(4) are shown on the right. The box plots show the distribution of contact numbers. The lower and upper bounds of the box are the first and the third quartiles respectively, while the line in middle is the median. The lower and upper limits of the dashed lines are the minimum and maximum number of contacts observed for cells for each combination of *α* and *D_T_*. The simulation was run for a total time of *T* = 1000 and updates in cell position were made every *δt* = 0.001.

We next investigated the effect of increasing diffusivity on the number of contacts for constant *α* (1, 10 and 20). Results from this set of simulations are shown in Figure 3. The red dotted line serves as a guide to the eye highlighting the trend observed. We see a steady increase in cell-cell contacts with diffusivity. Without diffusion, the test cell shows unidirectional motion towards the central cell and remains in contact throughout the simulation. Increasing diffusion increases the chance of test cell to go out of the radius of influence and come back again (trajectories 3 and 4).

**Figure 3.**
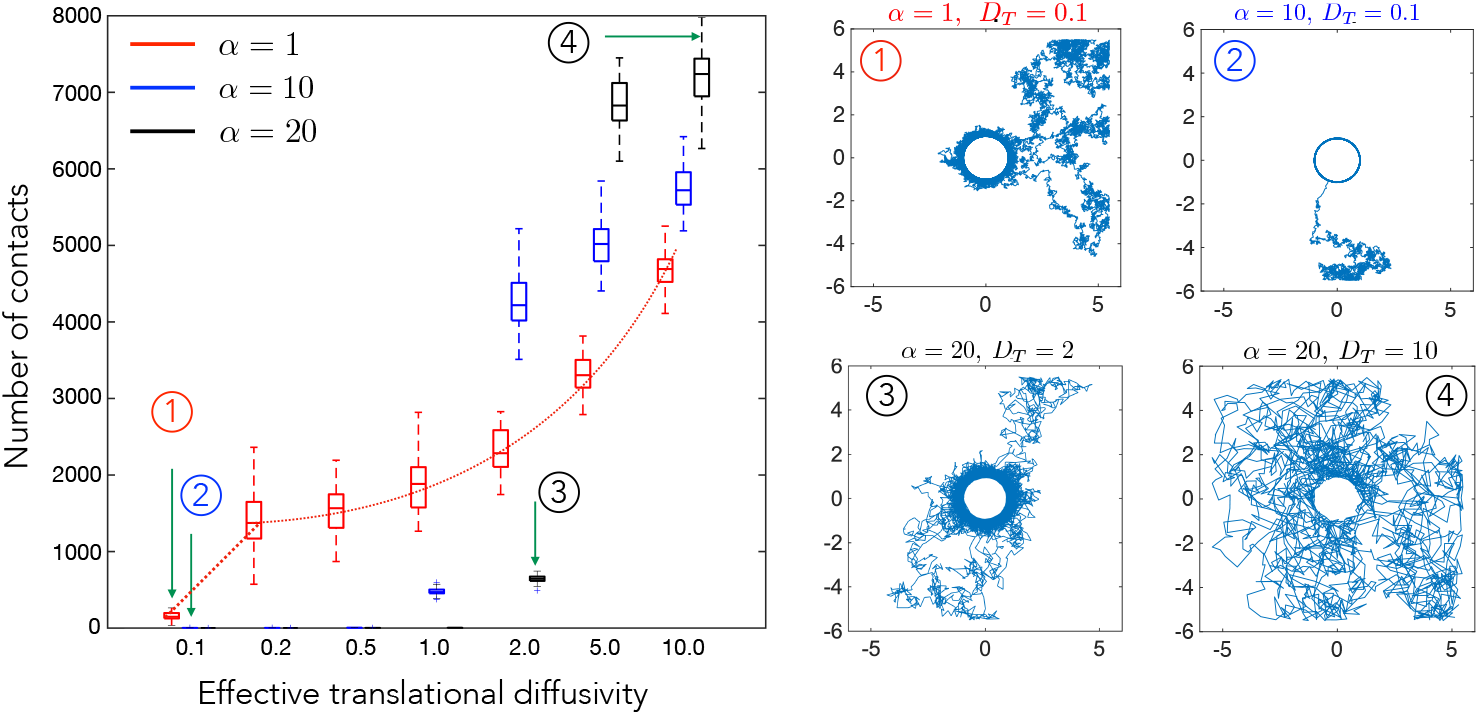
The number of cell–cell contact events in a fixed interval of time (*T* = 1000) plotted here as a function of the scaled effective diffusivity, *D_T_*, which represents the random motility of cell B. Here we show how the number of cell–cell contact varies for three different elastic interaction strength values, *α*, corresponding to substrates with three different stiffness. The highlighted points numbered from (1)-(4), show representative cell trajectories over long times and highlight how varying *α* and *D_T_* can yield states where the cells are in close proximity most of the time (low *D_T_*, high *α*) or states where cells interact rarely (high *D_T_*, low *α*). Interpretation of the box plots is the same as in Figure 2. The simulation was run for a total time of *T* = 1000 and updates in cell position were made every *δt* = 0.001.

Overall combining the results shown in Figures 2 and 3, we conclude that the number of contacts is maximized at an optimal value of the elastic interaction strength. If the elastic strength is too high or too low, the cell either makes stable contact or is too motile to make too many contacts. This optimal value scales with the diffusivity, which is a measure of the cell motility in our model.

### 4.2. Cell motility characteristics depend on elastic interactions

To quantify the long-time statistics of the motility of cell A in the elastic potential field generated by cell B, we analyze the mean squared displacement (MSD) as given by equation (9) from simulation. The metric MSD measured in terms of a delay time *τ* contains information about the short time mobility of a cell, the long time mobility of the cell, and additionally provides signatures of capture and trapping effects. Specifically, the slope of the mean square displacement can be used to extract effective exponents that provides insight on the relative importance of diffusion and elastic attractive interactions.

We plot the MSD in Fig. 4 for *D_T_* = 2 and *α* = 0.1, 1, 5, 10, 20,100. For *α* = 0.1, 1, 5,10, we find that the slope is close to 1, which suggests diffusion drives the motion of the cell and the attractive potential is not strong enough to influence the movement of the cell. For higher *α*, we observe a transition towards sub-diffusive behavior at *τ* ~ 0.5. At *α* = 20 (green line), the curve shows a significant decrease in slope at *τ* = 2, the time scale for which a test cell in average encounters the central cell for the first time and stays in contact for a while, as shown by trajectory 3, Figure 3. The slope then increases again, but remains less than 1 suggesting a sub-diffusive behavior in the long run. At *α* = 100 (blue line), the MSD saturates after initial diffusion to a zero slope which suggests that the motion is bounded, and it can only explore the circumference of the stationary cell.

**Figure 4.**
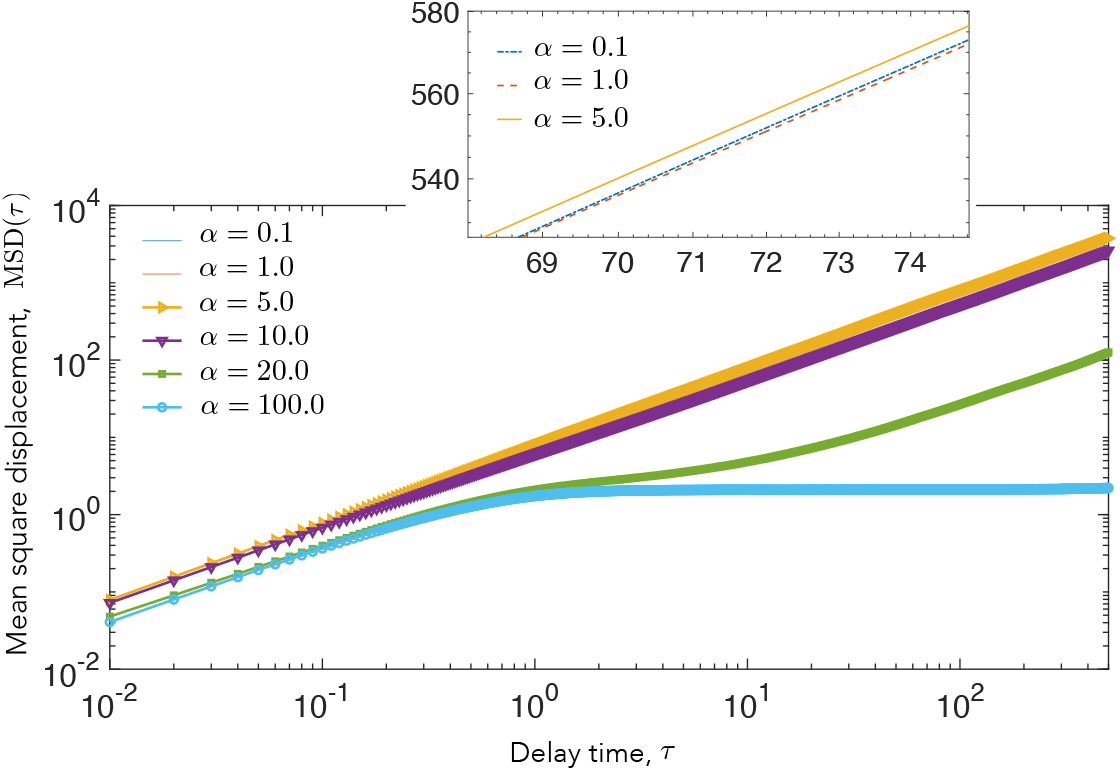
Mean square displacement. (MSD) as a function of the delay time interval *τ* (calculated from Equation 9), for the motile cell A is shown. Here we explore the variation in the MSD for various values of substrate-mediated elastic interactions, *α*. The diffusivity *D_T_* is held constant for these simulations with *D_T_* = 2. Other diffusivities were explored (results not shown). At low elastic interaction strengths, *α*, corresponding to stiff substrates, the cell shows a purely diffusive trajectory, whereas at higher values of *α*, the motile cell is captured by the strong attractive interaction from the stationary cell, resulting in a flattening of the MSD (blue curve). At an intermediate interaction regime (green curve), the motile cell makes repeated contact with the fixed cell but is never fully captured.

### 4.3. Elastic interactions lead to effective capture of motile cell

Taken together, our simulations suggest that strongly attractive elastic interactions can lead to stable contact between initially distant cells. We next explore the statistics of this “capture” process. Capture mechanisms underlying and influencing these statistics are potentially relevant for timescales of contact formation between initially well-separated motile cells that then form confluent monolayers, such as in mesenchymal–to–epithelial transitions during tissue morphogenesis [33].

Figures 2 and 3 suggest that the motile cell A (as it explores space and samples the potential field over its various trajectories) is attracted to the stationary cell with the attracting force increasing with decreasing distance *r*. Acting in tandem and superposed on this aspect of the motion is diffusion that allows A to wander away from B multiple times.

In order to understand how parameters *α* and *D_T_* affect this phenomenon, we tracked the number of cells inside the contact radius over the course of the simulation. The probability of cells inside the contact radius reached a steady state at time *t* < 100 for all parameters (Figure 5A). Keeping *α* constant and increasing *D_T_* the probability of cells being inside the contact radius decreases (Figure 5B). The steady-state probability *P_SS_* increases with increase in *α* for constant *D_T_* (Figure 5C). To understand the relationship between *P_SS_* and both *α* and *D_T_*, we investigated *P_SS_* for the ratio *α*/*D_T_* and showed that they remain constant for this ratio.

**Figure 5.**
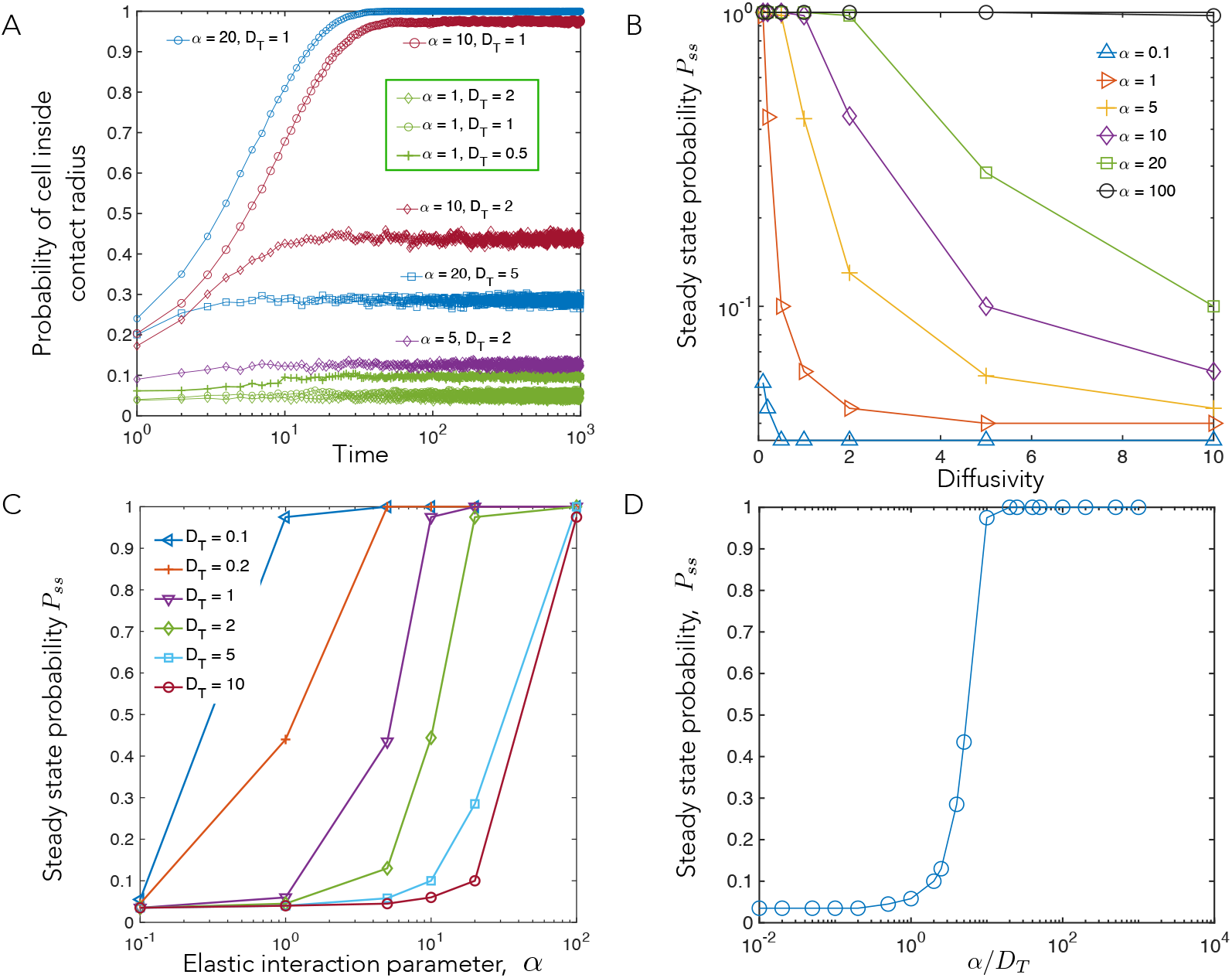
Capture statistics of motile cell. (A) Probability that cell B is inside contact radius as a function of time. (B and C) The dependence of steady state capture probability, *P_SS_*, *i.e*. the fraction of cells captured within the contact radius after a long time interval, on simulation parameters. (B) shows the dependence on diffusivity,*D_T_* at different values of the elastic interaction parameter, *α*, whereas (C) shows the dependence on *α* for different values of *D_T_*. (D) The steady state capture probability, *P_SS_*, data can be collapsed into a single master curve, when plotted vs. the key parameter, *alpha*/*D_T_*, the strength of the elastic interactions relative to the diffusivity. This is expected since our model steady state is a thermal equilibrium with effective temperature set by the noisy cell motility, *D_T_*, and the competition between attractive interactions and noise dictates the number of cells (cell trajectories) captured vs. the number that escape.

Plotting *P_SS_* vs *α*/*D_T_*, the strength of the elastic interactions relative to the diffusivity, we find that the data can be collapsed into a single master curve (Figure 5D). The collapse of our data and the master curve plotted in Figure 5D is expected since our model steady state is a thermal equilibrium with effective temperature set by the value of *D_T_*; the competition between attractive interactions and noise meanwhile dictates how many cells are captured vs. how many can escape.

This further justifies the notion introduced earlier of a *radius of influence*, that is, the distance from the stationary cell at which its elastic attractive tendency approximately balances the random noisy movements of the motile cell. Here we use a simple balance to estimate this radius of influence. Working in dimensionless units, we note that the dipolar interaction potential fall off as *α*/*r*^3^, while the effective temperature – a measure of the randomizing force – scales as *k_B_T* = *μ_T_D_T_*. Balancing these yields,

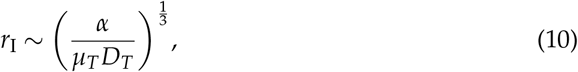

which explicitly shows the importance of the *α*/*D_T_* parameter. Thus, a stronger *α* from deformations exerted by the stationary cell (corresponding to softer substrate stiffness, or higher contractility) and lower random movements of the motile cell, *D_T_*, leads to a larger radius of influence. This in turn implies that the probability of being captured within the contact radius increases because the stationary cell can influence motile cells over a larger area.

### 4.4. Future work and perspectives: Anisotropic cell-cell elastic interactions

For polarized cells, that orient their cytoskeletal fibers and contractility along some principal axis, the cell-cell interaction potential is not isotropic. The individual cells on an elastic medium behave as force dipoles, with interaction potential energy having both attractive and repulsive regions that depend on mutual orientation of the two cells and their separation vector [19], as detailed in Appendix A. The force experienced by the motile cell has both radial and tangential components depending on its position and orientation relative to the central cell, and its direction is sensitive to the Poisson’s ratio of the elastic medium [34]. Thus, trajectories of cell A interacting with stationary cell B when the fully anisotropic interaction potential (Equation A1 and A2, Appendix A) is included will differ from trajectories observed in isotropic potentials. The difference arises in part due to an additional torque that reorients cell A to preferentially align with cell B as it moves towards it. Nonetheless, qualitative nature of the capture process and the observation of an effective region of influence will still remain valid.

To illustrate this we simulated the equilibrium orientation of uniformly spaced (pinned) test dipolar cells on a square lattice which are kept fixed in a square box of length 10*σ*. The Poisson’s ratio of the simulated substrate is 0.3 and *α* is 40. Results are shown in Figure 6. None of the cells overlap with the central stationary cell; they may rotate to reorient their dipole axis but are restricted from translating. We re-iterate that the cells on the lattice do not mutually interact with each other, but are only meant to illustrate the interaction of a test dipolar cell A placed at different spatial locations with the central stationary cell B. We note that fixed cells adjust the axis of their contractile dipoles in accordance to the potential field due to cell B (the dipole axis of B is fixed). Superposed on this are two trajectories corresponding to two cells that are freed from constraints and allowed to rotate and translate in response to the two-cell potential and thermal noise. The two cells start from their equilibrium orientation - i.e, they are first held pinned and allowed to reorient until the dipole axis attains a static value and then the pinning constraint is removed. Cells in the close vicinity of the central cell’s orientation axis exhibit a nearly linear motion to the pole of the fixed cell (trajectory in black). Cells away from the orientation axis take a longer route to come in contact with the central cell (trajectory in blue). The common attribute in both trajectories is that they prefer to adhere to the central cell’s pole, that is cell A as it moves towards B also continuously reorients in a manner that brings it into alignment with the cell B’s polar axis (the axis of the dipole).

**Figure 6.**
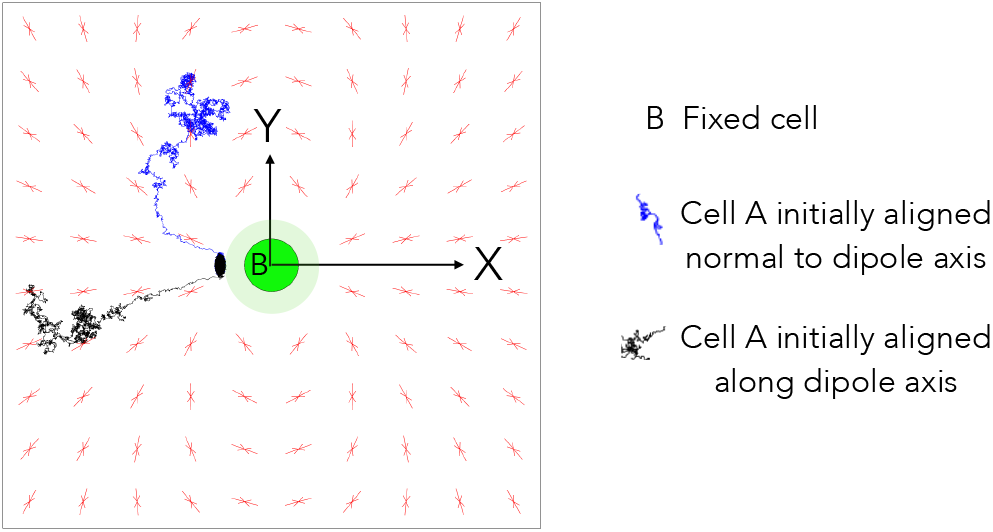
Dipolar cell orientation and trajectory. The equilibrium orientation of contractile cells fixed in position, but free to reorient, and that are uniformly distributed in a square box of size 10*σ*, are depicted by two arrows (red) pointing towards each other. Each cell is influenced by the central stationary cell B (green) and not by each other. Two possible trajectories of cell A (blue and black) are recorded for *D_T_* = 0.1, *α* = 40 for total time *T* = 500 with time steps of *dt* = 0.001. The cells did not have any self propulsion or rotational diffusion. The Poisson’s ratio *v* of the substrate was considered 0.3 for this simulation

## 5. Discussion

Using our model for cell contractility and motility, we computed several metrics of experimental relevance such as number of cell–cell contacts, the mean square displacement of a motile cell in the presence of elastic deformations induced by a cell in its vicinity, and associated capture statistics resulting from attractive interactions between two such cells. In each case, we predict how the computed metric depends on the elastic properties of the substrate, captured in the interaction parameter, *α* ~ 1/*E*, and on cell motility, captured by the effective diffusivity, *D_T_*.

Similar to the observations for pairs of endothelial cells mechanically interacting through the compliant substrates [3], we find that the motility and number of cell-cell contacts are lowered at large *α*, corresponding to softer substrates. This is because the elastic deformations of the substrate, and therefore, the cell–cell attractive interactions are stronger compared to the random motility. As observed in experiments, we also find that at intermediate interaction strength, the cells can make repeated contacts and withdrawals as shown in the contact number measurements. For very stiff substrates, that is low interaction strength, we find the cell remains diffusive and can migrate away from the stationary cell and does not make frequent contacts. Our findings would therefore suggest an optimal substrate stiffness at which contact frequency is maximal. These trends are also reflected in the MSD measurements. Unlike the experiment, we don’t find diffusive MSD for the strongly attractive case, but the MSD turns subdiffusive, suggesting perhaps that such high interaction strengths were not probed in experiment.

Biologically, such altered motility and contact formation could be relevant for forming stable adhesive contacts between cells and tissue development, including that of blood vessels during vasculogenesis [35]. We made several simplifying assumptions in the model (stated in section 2), including using a purely attractive and isotropic potential instead of the dipolar potential relevant for elongated and motile cells. Fig. 6 illustrates how the position and orientation of the motile cell with respect to the stationary cell leads to qualitatively different trajectories when the interaction potential is dipolar. Such an anisotropic potential is expected to lead to end–to–end alignment and contact formation of a pair of cells. With multiple cells, larger scale structures such as chains and networks of cells can result [19]. The influence of cellular motility on these structures will be the topic of a future study. The advantages of complementing experimental studies with modeling approaches as discussed in this paper is that hard to realize parameter regimes may be easily investigated. Furthermore, the role of different physical parameters may be clearly studied in isolation; a feature hard to achieve in an experimental setting.

In summary, our results illustrate how cell–cell mechanical interactions can lead to their mutual contact formation without requiring specific chemical factors to guide their motility, and how the substrate stiffness is an important control parameter in guiding cell motility and forming multi-cellular structures. The computational framework introduced and analyzed here can be extended to study durotaxis – that is, the modification of cell motility by variations in substrate elasticity at the single cell or tissue level and the motion of cells towards higher stiffness regions [36,37]. Understanding the mechanistic aspects of cell-cell interactions as done here has implications for regenerative medicine and tissue engineering and will guide and inform experiments exploring how cells communicate with each other in the process of organizing and moving collectively.

## Author Contributions

Conceptualization, K.D. and A.G.; methodology, A.G. and K.D.; software, A.G and S.B.; validation, S.B., K.D and A.G.; investigation, S.B.; resources, K.D. and A.G; writing, S.B., K.D and A.G. All authors have read and agreed to the published version of the manuscript.

## Institutional Review Board Statement

Not applicable.

## Data Availability Statement

Data is contained within the article or supplementary material.

## Abbreviations

The following abbreviations are used in this manuscript:

MSD: Mean Square Displacement

## Appendix A Model for a moving cell interacting with a stationary cell via substrate elasticity

The flat substrate is treated as being semi-infinite (Figure 1) and comprised of a linearly elastic, isotropic gel-like material with Young’s modulus *E* and Poisson’s ratio *v*, that capture its stiffness and compressibility respectively. The minimal model that describes the deformations created by cells exerting contractile forces on the substrate is a point-like force dipole [31]. Two identical dipolar cells denoted by *A* and *B* move in the upper plane (chosen to be the x-y plane, see Figure 1). Cell *A* is allowed to move and its dynamics is specified completely by its location on the substrate **r**^*A*^ (*t*) and by its self-propulsion direction **e**^*A*^ (*t*). Cell *B* is held fixed at point **r**^*B*^. As a result of the contractile dipoles exerted on the substrate the cells communicate elastically. The potential *W^AB^* characterizing this elastic interaction between the two cells is given by

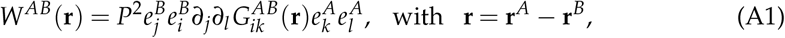

where *P* is the strength of the force dipole capturing the contractile stresses exerted by a cell on the medium. In writing (A1), we have made the plausible assumption that cells orient their cytoskeletal structures such as stress fibers and exert their traction primarily along their motility axis, such that the force dipole tensor, which captures the moment of their force distribution, is assumed to be, *P_ij_* = *Pe_i_e_j_*. The tensor

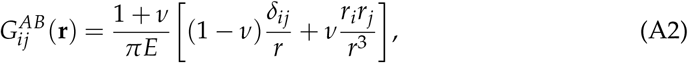

is the Green’s function that captures the displacement in the elastic medium at the location of one cell (dipole) caused by the application of a point force at the location of the other [38]. The partial derivatives in (A1) on the right hand side are taken with respect to relative position vector **r**. Standard Einstein notation has been chosen in writing the form of *W^AB^* and the derivatives in equations (A1) and (A2).

To obtain the force and torque balance equations that govern the dynamics of cell *A*, we make the simplifying assumption that the cells move in an overdamped fashion. This implies that hydrodynamic interactions between cells are ignored, and that each cell feels a resisting viscous frictional drag/torque that is proportional to its velocity/rotation rate. Conversely, when acted on by a force **F** or a torque **T**, a cell in this overdamped environment will move with velocity *μ_T_***F** or rotate at a rate *μ_R_***T** respectively. Here, *μ_T_* and *μ_R_* are appropriate mobility terms that depend on the cell size.

The micro-dynamics of cell *A* moving on the substrate is governed by the Langevin equations for the translation and rotary motion of cell. Recognizing that the elastic interaction generates (extra) forces and torques that act on each cell, and including the effects of fluctuating time dependent forces ***ζ***^*T*^(*t*) and torques ***ζ***^*R*^(*t*) originating from thermal noise, we can write the equations for the position and orientation of cell *A* in the presence of cell *B* as

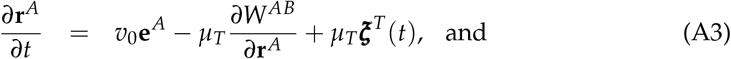

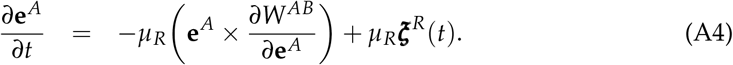

In an equilibrium situation, the random forces and torques are white noise terms and are related to one another by the equipartition and fluctuation-dissipation theorems: 〈**ζ**^*T*^(*t*)**ζ**^*T*^(*t*’)〉 = (2*k_B_T* / *μ_T_*)***δ****δ*(*t* – *t*’) where ***δ*** is the Kronecker delta function. For active cells however, these restrictions do not hold; these terms are set by active internal cell responses to the substrate properties. Equations (A1–A4) are used in the results illustrated in Figure 5.

In the bulk of the paper and for results presented in Figures 1–4, we use an isotropic version of the potential in equation (A1) that ignores orientational dynamics that are in general present for highly elongated cells. This assumes a separation of scales between the time over which cells reorient and the dipole axis changes and the time for the center of the cell to move significantly such as when the rotation noise in (A4) is significant. In this limit, one can average over the rapid reorientations of the cells and replace 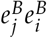 by *δ_ij_* and 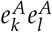 by *δ_kl_*. Equation (A1) then reduces to the simpler form that we employ in the main discussion of the paper and implement as a numerical simulation,

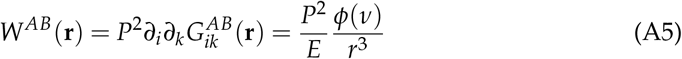

with the function *ϕ*(*v*) = (1 – *v*^2^)/*π* dependent solely on the Poisson ratio, and hence fixed in the simulation. Furthermore, since the dipole axis of cell *A* reorients in time scales much faster than its slower rate of translation, the *v_0_***e**^*A*^ term in (A3) simplifies to a time fluctuating variable with a mean that is roughly zero but with a non-zero variance. Thus its net effect may be incorporated by appropriately modifying the translational diffusivity. For an isotropic symmetric potential as here, the equation that needs to be solved is then

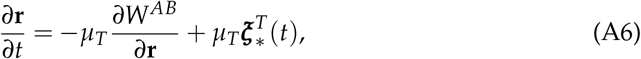

with the modified random force 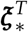 reflecting an effective translational diffusivity *D*_eff_ different from the thermal diffusivity *D*_0_, through a relation, 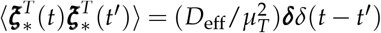. We define the dimensionless number *D_T_* ≡ *D*_eff_/*D*_0_. Consistent with this, we choose *μ_T_* = *D*_eff_/*k_B_T*.

